# HONeD-in on Brain Activity: Deconvolving Passive Diffusion on the Structural Network from Functional Brain Signals

**DOI:** 10.64898/2026.01.05.697753

**Authors:** Benjamin S Sipes, Fahimeh Arab, Srikantan S Nagarajan, Ashish Raj

## Abstract

Brain regions perform distinct computations, and their signals propagate through the whole-brain white matter network. Yet, mathematical models that describe this signal propagation via *purely passive diffusion* can predict a considerable amount of the observed functional connectivity between regions. This raises a critical question: if so much functional connectivity can be explained by a *passive* process, how can we isolate the *active* process? Here, we calculate in closed-form an estimate for such an active signal in functional MRI by spatially deconvolving the effect of passive signal spread over the brain’s structural connectivity using a higher-order network diffusion (HONeD) model. Across 770 Human Connectome Project subjects, we show that the resulting HONeD-innovation (HONeD-in) signal 1) sparsifies functional connectivity while retaining a well-connected network, 2) remodels resting-state networks (RSNs), 3) mixes the unimodal– multimodal hierarchical organization of RSNs into a circle with no clear hierarchy, and 4) deblurs task-activation maps. Together, our results highlight HONeD deconvolution as a generalizable new way to study resting-state and task fMRI brain signals.

## Introduction

Neurons connected in a network integrate and transmit the signals they receive. When averaged over large populations, these complex nonlinear dynamics blur into macroscopic signals that are often well approximated by linear models [48]. This macroscopic signal is akin to a fluid traveling through the structural pathways between brain regions. Yet, like a fluid, we may expect that if a brain region exhibits high activity (i.e., a high ‘concentration’ of signal), then that signal should passively ‘flow’ down a concentration gradient to connected regions with less signal. This is the well-known physical process of passive diffusion. Therefore, in macroscopic brain imaging signals, we may consider two coincident types of signal movement through white matter: (1) signal moving according to *purely passive network diffusion*, and (2) signals that *cannot* be predicted by passive diffusion, which could be interpreted as *actively generated and transmitted signals*. Is it possible that our attempts to study brain activity and connectivity with macroscopic neuroimaging are in fact confounded by an ‘undercurrent’ of passive network diffusion?

The presence of such a passive diffusion-based signal in macroscopic neuroimaging has been reported in fMRI, where a significant amount of the brain’s functional connectivity (FC) can be predicted from passive diffusion along structural connectivity (SC) [1, 2]. In magnetoencephalography (MEG), related models based on network diffusion over SC are capable of predicting the brain’s frequency power spectrum [53, 71]. Related ideas appear in linear time-invariant (LTI) state-space models commonly used in network control theory, which explicitly model intrinsic linear dynamics over SC separate from external (‘exogenous’) system inputs [29–31, 38, 6]. In other signal processing contexts, such a component unexplained by the model’s intrinsic dynamics has been termed an “*innovation*” signal [45]. However, if passive network diffusion can explain much of the macroscopic brain signal we measure, then it may be contributing substantially to canonical findings in neuroscience, including resting-state networks and task-related activations. To what extent, then, are these phenomena driven by passive diffusion rather than an active innovation signal?

The endeavor to recover these active sources of brain signal has been a long-standing pursuit in neuroscience. Early on, independent component analysis (ICA) was proposed as a method for blind source separation to recover the independent sources of brain activity [12]. ICA and other techniques, such as von Mises-Fisher clustering [74] and InfoMap [56], have been used to robustly identify whole-brain patterns of co-activation, commonly referred to as resting-state networks (RSNs), which have been treated as intrinsic network-level sources in the brain. These RSNs show remarkable stability across varying analytical techniques [60, 59] and neuroimaging modalities [13, 11, 76]. Interestingly, several studies indicate that canonical RSNs are related to and can emerge from the topology of SC itself [3, 2, 73, 62], raising the possibility that a process over SC, such as passive diffusion, lies at the heart of RSN organization. Other prior work has tried recovering innovation signals in the brain using spatio-temporal hemodynamic response function (HRF) priors (e.g., “Total Activation” [37]) or dynamical modeling with Kalman filtering [32]. Overall, the goal to recover the brain’s innovation signal is critical since our interpretations of both resting-state and task-based brain activity depend on the assumption that our neuroimaging measurements relate to what the brain is *actively* doing.

However, previous approaches to recover active innovation signals in the brain are significantly limited since they lack a physics-based model on SC. Without such a model, deriving an innovation signal is not only ill-posed, with many possible parameterizations and solutions, but also divorced from the underlying biology of white matter connectivity. What is needed is a model for passive network diffusion over SC that can be inverted in such a way that performs a *graph deconvolution*, thereby recovering the brain signals that are not blurred out by spreading over the network [20, 75, 61]. Such a graph deconvolution, grounded in the physics of passive diffusion on SC, provides a biological interpretation for the recovered innovation signal: it corresponds to a mixture of computations occurring within regions and to actively transmitted signals shared between regions. A subtle but crucial point is that the recovered innovation signal can *still be constrained by the underlying SC* since the graph deconvolution *amplifies* the contribution of SC modes that *are not* related to passive signal spread governed by the physics-based model.

Here, we propose such a graph deconvolution approach using a physics-based higher-order network diffusion (HONeD) model to predict the functional MRI (fMRI) signal component attributable to passive network diffusion over SC. As we shall show below, the HONeD model fits better to empirical data, and its use prevents exponential amplification of high-frequency SC modes that arise when inverting simpler diffusion kernels. We used HONeD to deconvolve our model prediction in *closed-form* to recover the HONeD-innovation (HONeD-in) signal underlying brain activity. In the absence of ground truth, we thoroughly validate our HONeD deconvolution approach in simulation, showing that we can accurately recover ≈ 80% of the original innovation signal with robustness to noise. In empirical fMRI data, we show that HONeD-in signals (1) sparsify FC and change its network topology; (2) fracture conventional RSNs into decoupled and interdigitated networks; (3) reorganize traditional RSN hierarchies in FC gradients, transforming the unimodal–multimodal divide into an organization without clear stratification; and (4) yield deblurred activation maps in task-based general linear model (GLM) analyses. These findings together suggest that dominant network phenomena in fMRI may be deeply related to the brain’s SC. HONeD deconvolution thus offers a fundamentally new technique to investigate brain activity in neuroimaging and ensure that our findings are not confounded by passive diffusion over structure.

## Methods

### Theory

#### General Linear Time-Invariant Model for Brain Activity

In the study of Linear Time-Invariant (LTI) systems, a general state-space equation for how a signal behaves in a system can be written as:

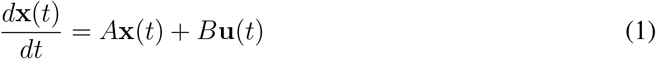

Here, **x**(*t*) ∈ ℝ^*n*^ is a vector specifying the state of the system’s signal across *n* nodes, *A* ∈ ℝ^*n*×*n*^ is a state transition matrix (i.e., the system’s “internal dynamics”), **u**(*t*) ∈ ℝ^*m*^ is a vector specifying the exogenous input into the system at *m* inputs, and *B* ∈ ℝ^*n*×*m*^ is an actuation matrix that specifies how the input distributes through the system. In neuroscience, this equation is commonly invoked when studying network control theory, where the internal system dynamics *A* can be taken as the structural connectivity (SC), and certain aggregate properties of *B* and/or **u**(*t*) are estimated [29, 38].

Here, we leverage this framework to model intrinsic brain dynamics as diffusion over structural connectivity.

### The Graph Laplacian and Higher-Order Network Diffusion

We construct the state matrix *A* using the *graph Laplacian*, making *A* a physics-based operator for network diffusion. A full first-principles derivation and explication for the relationship between the graph Laplacian and the physics of signal diffusion has been well documented [49, 1]. For completeness, we will provide an abbreviated description here.

Consider the brain’s structural connectivity matrix (*C* ∈ ℝ^*n*×*n*^), where each row/column represents a single brain region of interest (ROI) and the matrix entry (*C*_*ij*_) corresponds to the connectivity strength between regions *i* and *j*. Then the way in which a signal can be expected to change under passive diffusion is 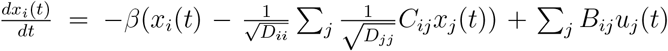, where *D*_*ii*_ = ∑_*j*_ *C*_*ij*_ corresponds to the amount of connectivity (degree) at a given region *i*. Vectorizing this equation reveals the degree normalized graph Laplacian:

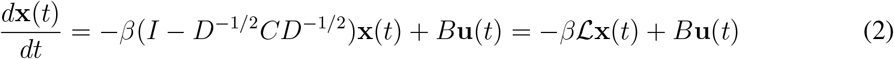

The graph Laplacian (ℒ ∈ ℝ^*n*×*n*^) is a graph analogue to the Laplace-Beltrami operator for manifolds [36], and it is deeply related to the physics of diffusion over networks [49].

However, note that the above case is in fact a specific linearized truncation of a more general way to specify diffusion through a network. Instead of considering merely −*β*ℒ as the operator for diffusion, we can generalize this operator to be of the form

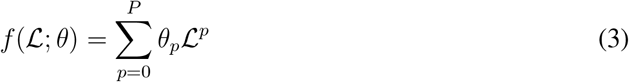

Here, *P* is the order of the Taylor series approximation for a generalized “higher-order network diffusion” (HONeD) operator, and *θ* ∈ ℝ^*P* +1^ is the vector of HONeD parameters. A key advantage of this generalized form is that it accounts for multi-step pathways through the brain’s structural network^1^ [5, 14].

It is relevant to note that being diagonalizable (ℒ = *U* Λ*U* ^⊤^) with orthogonal eigenvectors (hereafter called “harmonics”) *U* and eigenvalues Λ means that any polynomial function of this matrix can be written as a function of its eigenvalues^2^:

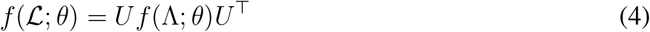

The graph Laplacian harmonics can be interpreted as standing waves sustained on the structure of the network. Decomposing the Laplacian into its harmonics provides an intuitive and computationally useful way to fit the parameters (*θ*_*p*_) of the HONeD operator, which we’ll describe in detail below.

With our HONeD operator in hand, let’s return to the LTI model and derive the HONeD-innovation (HONeD-in) signal. Henceforth, we specify *A* = *f* (ℒ; *θ*) such that the internal system dynamics is governed by the physics-based HONeD diffusion operator over SC.

### Deriving the Innovation Signal

In this work, we seek to estimate a signal **z**(*t*) = *B***u**(*t*), since *B* and **u**(*t*) are not individually identifiable, and therefore our estimate bundles together the exogenous system input **u** with the system actuation *B*. Hence, our differential equation takes the form:

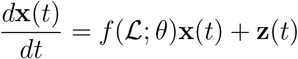

The solution to this inhomogeneous ODE is:

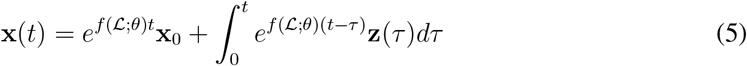

Here, we assume that the innovation signal behaves like a Dirac delta function of the form **z**(*t*) = ∑_*k*_ **z**_*k*_*δ*(*t* − *t*_*k*_) with *t*_*k*_ = *k*Δ*t*, which is a sparse signal in time, and where we assume the convention where each pulse is the beginning of each TR. Assuming the innovation signal behaves like a delta function is aligned with previous work on estimating innovation signals in fMRI [37]^3^. Using this formulation, we show that we can calculate the HONeD-in signal in discrete time as:

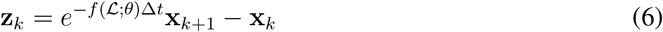

For a thorough mathematical derivation, see the Supplemental Material Section 1. However, this closed-form equation requires knowing the correct *parameterization* (*θ*) for HONeD *f* ( ; *θ*). Hence, we next show how we estimate the HONeD parameters under a homogeneous passive diffusion model.

### Estimating HONeD Parameters

To approximate the effect of homogeneous passive diffusion, we take the general LTI equation, but without any specific exogenous drive: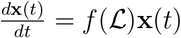, which has the solution **x**(*t*) = *e*^*f*(ℒ)*t*^**x**_0_ = *Ue*^*f*(Λ)*t*^*U* ^⊤^**x**_0_. ^4^ Importantly, this model makes a different assumption about the initial condition of the system–the homogeneous HONeD model treats the “initial condition” as an independent and identically distributed (i.i.d.) Gaussian random variable such that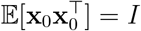. ^5^

We use this homogeneous HONeD model to predict the covariance of fMRI signals during resting-state:

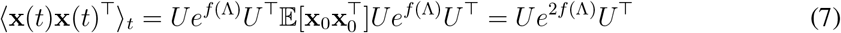

We wrote this model prediction in terms of the Laplacian’s harmonic decomposition to highlight how HONeD also *predicts the shape of harmonic power*. Specifically, let us define 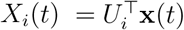 as the projection of the time series to the space of the *i*^th^ graph harmonic (Figure 1a) and graph harmonic power as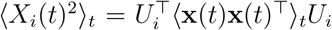. Then our model, equation (7), can be evaluated as a scalar function:

**Figure 1:**
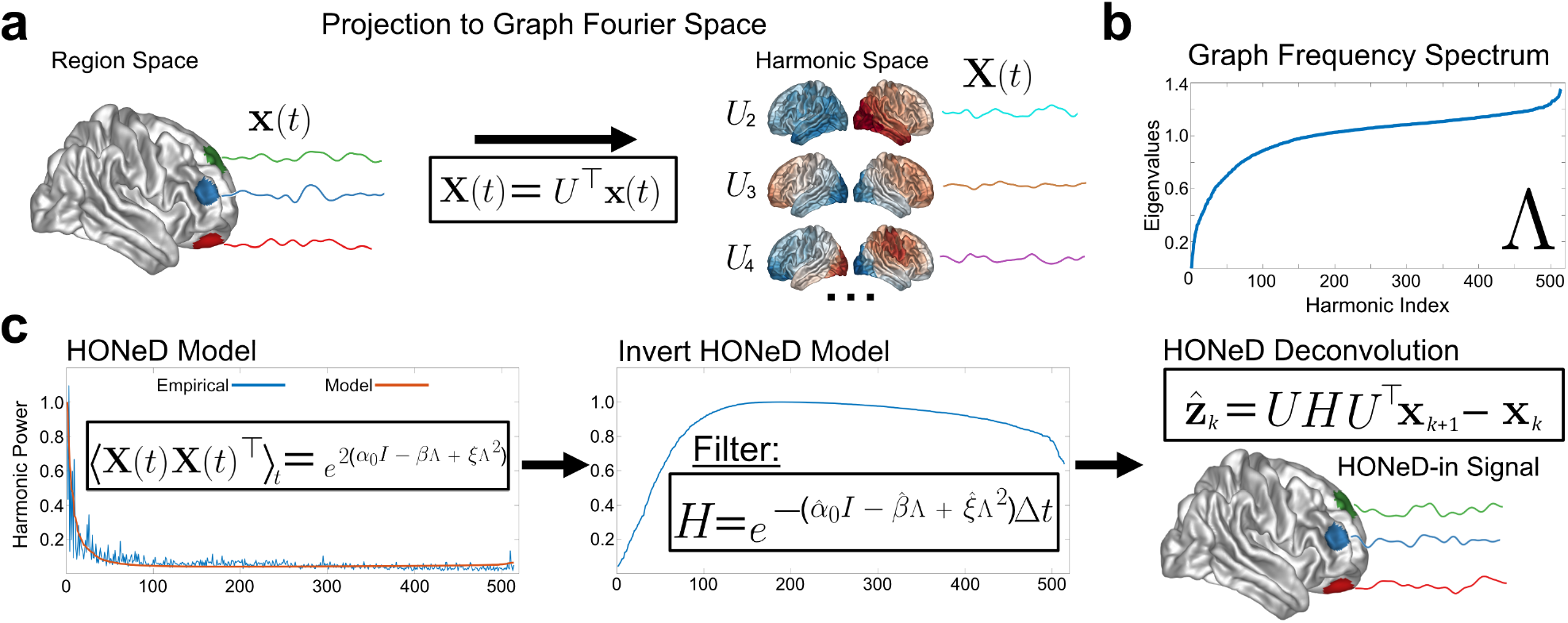
(a) A depiction of the Graph Fourier Transform taking signals in region space and projecting them into the harmonic space. (b) The graph frequency spectrum (i.e., Laplacian eigenvalues), which are used to compute the HONeD model prediction. (c) The HONeD model fits to the empirical harmonic signal power. This prediction is inverted to create the HONeD filter which is subsequently applied to the original time series to obtain the estimated HONeD-innovation signal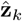. (Notation: bold denotes time series signals, lower-case denotes region space signals, and upper-case denotes harmonic space signals.)

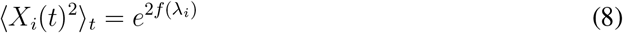

To fit HONeD to empirical fMRI, we solve a constrained least-squares regression problem on the log-transformed harmonic power: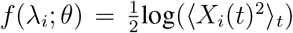. Since least-squares fitting to polynomials is a convex quadratic optimization problem with globally optimal solutions, this model formulation is both fast to solve and comes with minima guarantees [8]. Additionally, we fit the HONeD model with a constraint that the polynomial be non-decreasing at the largest eigenvalue (*λ*_max_), meaning that its derivative is non-negative. This constraint ensures that inverting the HONeD kernel does not overly amplify the highest eigenvalue harmonics. Thus, we use the following optimization function to estimate the subject-specific HONeD parameters 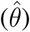 in harmonic space:

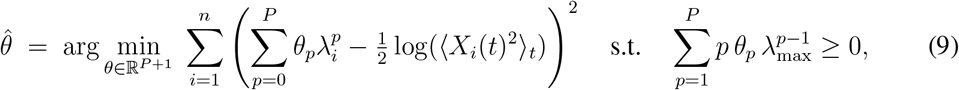

So far, we have kept the form of HONeD general, but in practice we must choose a polynomial order. For the current work, we chose to limit HONeD to two steps (*P* = 2) through the network, since previous work has shown that two steps through SC nearly saturate SC’s ability to predict functional connectivity [5], and we verified that two-step diffusion was also optimal for HONeD in this dataset (see Supplemental Figure 7). Therefore, the HONeD operator for the present work becomes

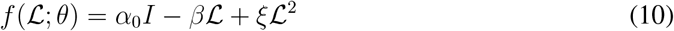

We denote the terms from the Taylor expansion *θ*_0_ = *α*_0_, *θ*_1_ = −*β*, and *θ*_2_ = *ξ* for ease of referencing. Since empirical work has shown that *θ*_1_ *<* 0, we reference *θ*_1_ = −*β* to make *θ*_1_’s empirical negativity clear and to align terminology with previous network diffusion model work [1, 2, 51]. By fitting the HONeD model to the empirical harmonic power, we estimate subjectspecific parameters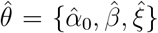, which we describe in detail below. An overview of the study theory is provided in Figure 1.

### Estimating the HONeD-in signal

Having inferred the parameters of the homogeneous HONeD model, we finally estimate the HONeD-innovation (HONeD-in) signal as

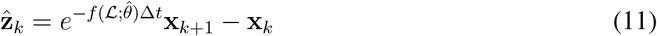

In words, at each fMRI sample, the HONeD-in signal is obtained by “unwinding” the system defined by our HONeD operator, then comparing with the previous state. As such, 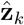 is estimated as a *graph-deconvolution*.

### Dataset and Processing

We used 770 subjects from the Human Connectome Project (HCP) Young Adult dataset in this study [69] ^6^. We included all HCP subjects that had both diffusion and functional MRI, and excluded subjects with quality control issues. Unprocessed HCP data were downloaded and organized into BIDS format. Anatomical scans were collected with an MPRAGE T1-weighted sequence. Functional MRI (fMRI) data included four 15-minute rest sessions and two task sessions for six tasks (Motor, Language, Gambling, Social, Relational, and Working Memory), all with a repetition time (TR) of 720 ms. Additionally, we used the pre-processed diffusion tensor imaging (DTI) data made available by HCP, which includes merging high fidelity directions across multiple multi-shell DTI acquisitions (b = 1000, 2000, & 3000) and correcting for phase-encoding polarity distortion [25]. We applied a state-of-the-art image processing pipeline, micapipe [15], which processes the anatomical, functional, and diffusion MRI data in a coherent framework to produce subject-specific structural connectivity and functional regional time series in the Schaefer atlas with 500 cortical regions [59], each with connectivity to an additional 14 subcortical regions [50].

The structural connectivity was computed using MRtrix3 and probabilistic tractography, generating 10 million streamlines throughout the gray matter white matter interface (maximum tract length = 400, minimum length = 10, cutoff = 0.06, step = 0.5, angle curvature = 22.5°) using the iFOD2 algorithm and 3-tissue anatomically constrained tractography [15]. Tractograms were filtered using SIFT2 [63], reweighting streamlines by cross-sectional multipliers to provide connection densities that are biologically valid measures of fiber connectivity.

The fMRI underwent image re-orientation, motion and distortion correction, and nuisance signal regression (white matter, cerebrospinal fluid (CSF), and frame-wise displacement spikes). Volumetric time series were mapped to the native FreeSurfer space using boundary-based registration [27]. Vertex-level time series were averaged into parcels defined by the Schaefer atlas with 500 cortical regions [59]. Parcel-level time series were bandpass filtered to be less than 0.1Hz, then underwent global signal regression (GSR). We computed the FC as the Fisher Z-transform of Pearson correlations between parcel-level time series [21, 68].

### Simulation Analysis

We evaluated our HONeD deconvolution method by creating a “ground-truth” innovation signal by randomly permuting region order in empirical fMRI time series following GSR. Since HONeD is based on propagation over SC, we randomly shuffle fMRI time series to destroy its spatial correspondence with SC but retain all natural fMRI statistics and correlations between signals, making it a structured innovation signal. Then, we run this innovation signal through the LTI equation using the HONeD operator, equation (10), and Euler integration to generate a simulated “natural” signal 1000 times. Since network diffusion imposes a new global signal^7^, we perform GSR on the simulated “natural” signal and then fit our HONeD model to the post-GSR “natural” time series to estimate the innovation signal.

We determined how well HONeD deconvolution recovered the ground-truth innovation signal by computing two complementary metrics of signal recovery, normalized by the HONeD simulated “natural” signal. In the following, 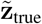 is the ground-truth innovation signal entered into the HONeD ODE, 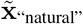 is the simulated signal after running the ground truth signal through the HONeD ODE, and 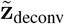 is the estimated deconvolved innovation signal.

First, we computed a normalized mean squared error (MSE) as:

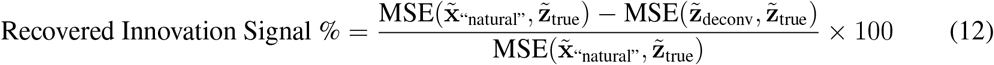

Second, we computed an analogous measure for FC:

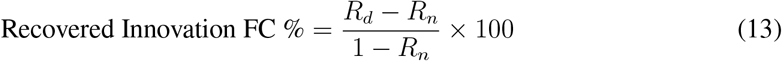

where *R*_*d*_ = corr(FC_true_, FC_deconv_) and *R*_*n*_ = corr(FC_true_, FC_”natural”_), where again FC_true_ is the functional connectivity computed from the ground-truth innovation signal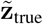. We also evaluated how well HONeD parameters were recovered and how optimal HONeD deconvolution behaves around the ground-truth parameters.

We leveraged this simulation to thoroughly evaluate the performance of HONeD deconvolution across a variety of conditions. To assess the optimal performance, we used the subject-specific SC in both the simulation and the HONeD filter estimation. To test generalizability of HONeD in cases where subject-specific SC may not be available, we estimated the HONeD filter using a consensus SC (i.e., mean across all 770 HCP subjects). To evaluate HONeD deconvolution’s robustness to SC estimation error, we added random Rician noise to subject-specific SC. We scaled the standard deviation of the Rician distribution to the standard deviation of the original SC edge weight distribution. We evaluated Rician noise at the following scales: *σ* = {0.25, 0.5, 1, 2} (e.g., when *σ* = 1, the Rician standard deviation matches the empirical SC’s standard deviation). Similarly, we evaluated HONeD deconvolution’s robustness to additive Gaussian noise by first z-scoring the HONeD-generated “natural” fMRI signal then adding Gaussian noise with the same noise scales: *σ* = {0.25, 0.5, 1, 2}. We also evaluated how HONeD deconvolution performed using a randomized SC, which rewired empirical SC edges at least 5 times, only preserving the network’s binary degree distribution and connectedness [57]. Since HONeD introduces a global signal into the data, meaning also that HONeD deconvolution performs a type of GSR, we compared the ground truth innovation signal and FC to the simulated post-GSR “natural” signal, thereby testing whether GSR recovers any innovation signal or FC structure.

### Graph Theoretical Analyses

Recall that HONeD predicts the amount of fMRI signal governed by passive diffusion driven by i.i.d. Gaussian white noise. If passive diffusion with uniform actuation (*B* = *I*) is all there is to brain connectivity, the HONeD-in signal should have an FC that approximates the identity matrix with no structure in its network connectivity. Therefore, to evaluate whether the HONeD-in signal *does* in fact have meaningful network structure, we analyzed the graph theoretical properties of the “HONeD-in FC” and compared them to the same graph metrics from the post-GSR “natural FC,” as well as network randomized versions of the empirical networks and a network composed of only Gaussian noise. To align with traditional network neuroscience approaches, we computed the Pearson correlation between each region’s time series, then performed the Fisher Z-transform to obtain FC [21, 68]. For all graph theoretical analyses, we used only the positive FC edges, discarding all negative connections.

We compared natural to HONeD-in FC across traditional graph metric properties using the brain connectivity toolbox [57]. These metrics were modularity, mean clustering coefficient, mean betweenness centrality, mean eccentricity, the rich club (area under curve), and characteristic path length (CPL). These measures index a wide array of graph topological properties, especially the integrative and segregative network topology. Broadly, rich club AUC, betweenness centrality, and clustering coefficient indicate more integration while modularity, eccentricity, and CPL indicate more segregation [16]. Since HONeD deconvolution suppresses the integrative harmonics [62], we predicted that HONeD-in FC should show decreases in measures of network integration, resulting in a sparse network.

### Resting-State Network and FC-Gradient Analyses

Functional resting-state networks (RSNs) have become a cornerstone of neuroscience research. RSNs have previously been identified using von Mises-Fisher clustering on consensus FC [74, 59]. Therefore, we also evaluated the effect of HONeD deconvolution on the identification of these networks. We replicated this previous approach by performing von Mises-Fisher clustering with 7 clusters on the natural and HONeD-in consensus FCs with only positive edges across 10,000 initializations [33, 4, 39]. In this study, we are using the HCP dataset, which was the original dataset used to identify these networks in the Schaefer atlas [59]. Although the pre-processing differences may result in variation, we expect the natural FCs to align well with the canonical organization. We assessed the clustering performance with cross-validation in the same fashion as Yeo et al. [74]. That is, we identified RSNs using randomly selected half of the brain’s regions, then we predicted the RSN assignment using the Euclidean distance to the nearest centroid. We re-ran this cross-validation 1000 times for each clustering order (*K*). This yielded an accuracy distribution which we used as our measure of clustering stability.

Moreover, there has been much work investigating the properties of “functional gradients,” which are the Laplacian diffusion map eigenvectors of FC originally described by Margulies et al. [44]. This seminal work showed that these gradients intimately relate to RSNs: the first principle gradient organized a multimodal to unimodal hierarchy from the default-mode network (DMN) to visual/motor areas; the second gradient further separated the visual and motor cortices. To assess whether HONeD-in signals alter this hierarchy, we computed two FC-gradients for both the natural and HONeD-in consensus FC and evaluated whether FC-gradients were left unchanged by removing the effect of passive signal diffusion through SC. Then, we investigated how brain regions and RSNs derived from the HONeD-in FC organized around HONeD-in FC Gradients.

### Task-Contrast Analysis

We assessed how HONeD deconvolution influenced the activity across different HCP tasks. We followed a traditional approach taken in task fMRI studies by convolving a canonical hemodynamic response function (HRF) with the time-wise binary vector indicating task-blocks of interest. The form of the HRF was estimated for each subject directly from the data using the resting-state time series using the rsHRF package [72]. Since HONeD deconvolution may change the shape of the HRF, we estimated these HRFs separately for the natural and HONeD-in signals. We then used a general linear model (GLM) to compute the association between each parceled region’s time series and the HRF-convolved response time series with 1-step autoregressive prewhitening, which controls for temporal autocorrelation [23]. We computed region-wise contrast in the regression weights from the GLM for each subject as a first-level analysis (i.e., Δ*b* = *b*_task_ − *b*_rest_) ^8^, followed by a second-level analysis as a univariate t-test across all beta weight contrasts across subjects, using Bonferroni correction for multiple comparisons [17]. This produced test-statistic maps that we used to evaluate the differences in “regional activity” between the natural and HONeD-in signals. Note that, unlike traditional fMRI task-based analyses, we assessed the task activations using the post-GSR signal to ensure an appropriate comparison with other results.

## Results

### HONeD Fitting Performance

We found that an order-2 HONeD operator provided the best improvement on harmonic prediction given our model constraints (Supplemental Figure 7). The homogeneous HONeD model fit well to the empirical harmonic power across subjects in both mean squared error (MSE = 0.14(±0.03)) and Pearson correlation (R = 0.85(±0.02)). HONeD passive diffusion parameters across all 770 HCP subjects were *β* = 2.98(±0.26) and *ξ* = 1.07(±0.11) (Supplemental Figure 8).

### HONeD Deconvolution Simulation

We found that HONeD deconvolution was well-posed under simulations following the HONeD model. Our method for estimating HONeD parameters predicts close to the ground-truth simulated parameters on average (True *β* = 3, Estimated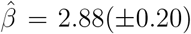; True *ξ* = 1.2, Estimated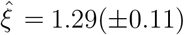; Figure 2a & b). We also evaluated whether the true parameters were indeed the optimal parameters for the deconvolution. We found that there was a range of ideal deconvolution parameters, and that the true parameters were indeed near the center of this range (Figure 2c). Since *β* and *ξ* can create similar deconvolution filter shapes when they change together, it makes sense that a range of correlated parameters can identify equally good deconvolution estimates.

**Figure 2:**
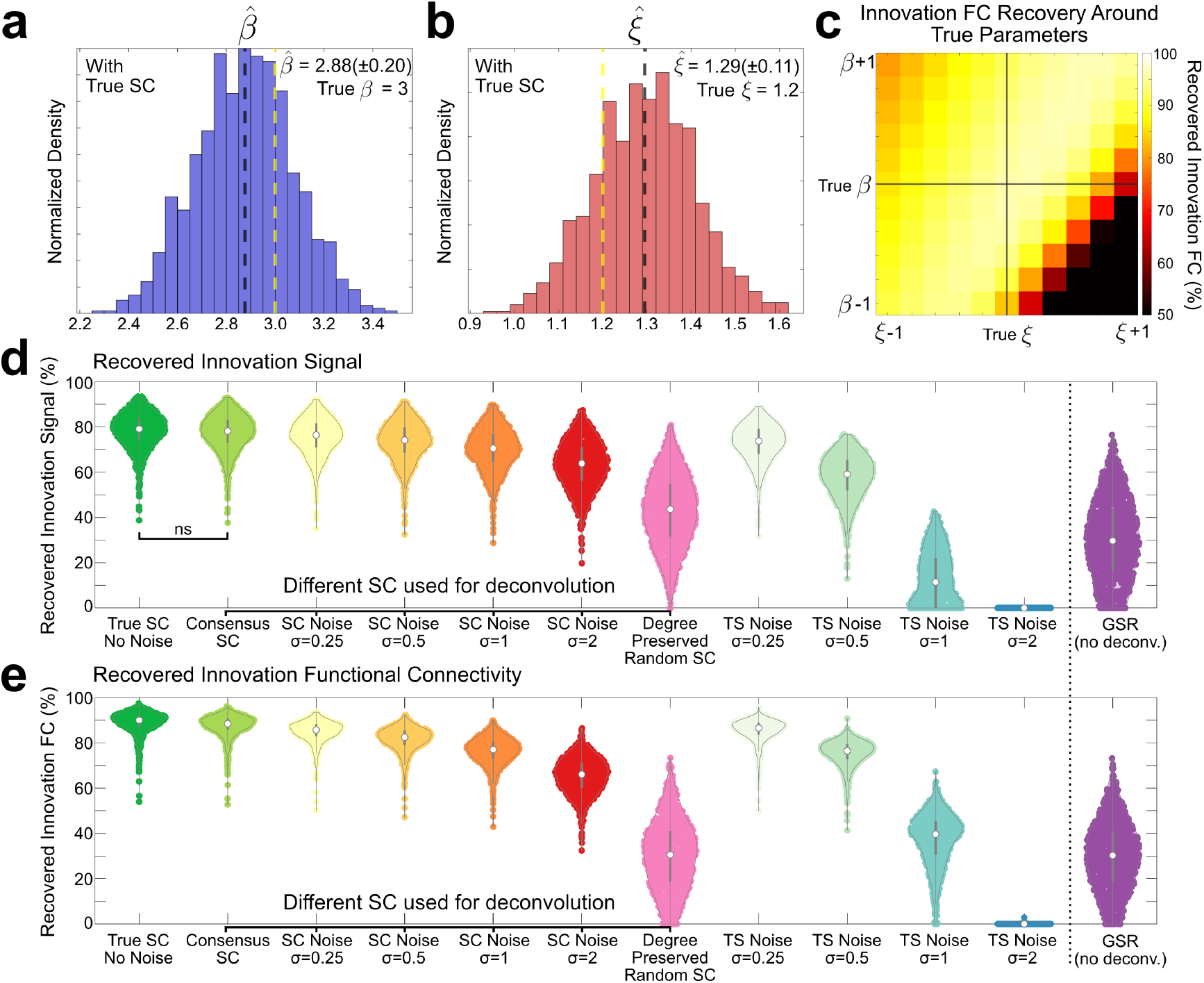
HONeD Simulation Results. (a & b) The distributions of estimated 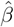 and 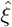. The dashed black line indicates the sample mean while the dashed yellow line indicates the simulated ground-truth value. (c) A heatmap showing how variations around the ground truth parameters influence the recovery between ground truth innovation FC and the deconvolved innovation FC. (d) Violin plots showing the Recovered Innovation Signal (%) across different simulation conditions, and (e) shows the Recovered Innovation FC (%). For visualization, we set all negative recovery values to zero. Both showcase HONeD deconvolution robustness to noise and clear improvement over GSR alone. We labeled all pair-wise comparisons that were non-significant (ns) under a Student’s t-test. Where violins are not labeled, pairwise distributions are significantly different after Bonferroni correction.

We next evaluated how well HONeD deconvolution could recover the original time series and FC across a range of conditions (Figure 2d & e). Unsurprisingly, subject-specific SC with no noise performed best, recovering 78% of the innovation time series and 89% of the FC. Using the consensus structural connectivity (SC) performed equally well for recovering the innovation signal, and slightly worse for FC (≈88%). More impressively, HONeD deconvolution was highly robust to SC noise, still recovering 63% of the innovation signal and 65% of the innovation FC even when the SC edge-wise noise was double the network’s standard deviation. The degree-preserving randomization, as expected, fared worse than noise, recovering only 43% of the innovation signal and 30% of the innovation FC. HONeD deconvolution was moderately robust to additive white noise: noise at half the signal’s standard deviation still recovered 58% of the innovation signal and 75% of the FC. The innovation signal was not recoverable when time series noise was twice the signal’s standard deviation. We finally evaluated how well only performing global signal regression (GSR) contributed to recovering the innovation signal. We found that GSR only recovered 30% of both the ground truth innovation signal the ground truth innovation FC.

Together, these results affirm that fitting HONeD can closely estimate the diffusion coefficients arising from signal propagation through SC, with robustness to noise sources, and significant recovery of underlying innovation signals and their covariance.

### HONeD-in FC is a Sparse Connected Network

After computing the HONeD-in signal, we computed its FC and compared it with “natural” FC (post-GSR) across several graph theoretical metrics. First, we found that HONeD-in FC significantly restructured the FC, increasing connections between canonical resting-state networks (RSNs) while decreasing connections within RSNs (Figure 3a & b; edges thresholded by Bonferroni correction). We next compared the consensus FCs (shown in Figure 3a) by their degree and edge distributions. We found that HONeD-in FC significantly *decreased* the overall degree of all nodes, and resulted in a narrower distribution with a heavier high degree tail (Figure 3c, left). Conversely, HONeD-in FC had a significant, but subtle, *increased* edge distribution with most edges being nearly zero, and a sparse set of non-zero edges (Figure 3c, middle). In an edge-to-edge comparison, we observe that the effect of deconvolution is to make negative edges closer to zero while acting as a soft-threshold to many high-strength edges, even converting some moderate FC connections to negative connections (Figure 3c, right).

**Figure 3:**
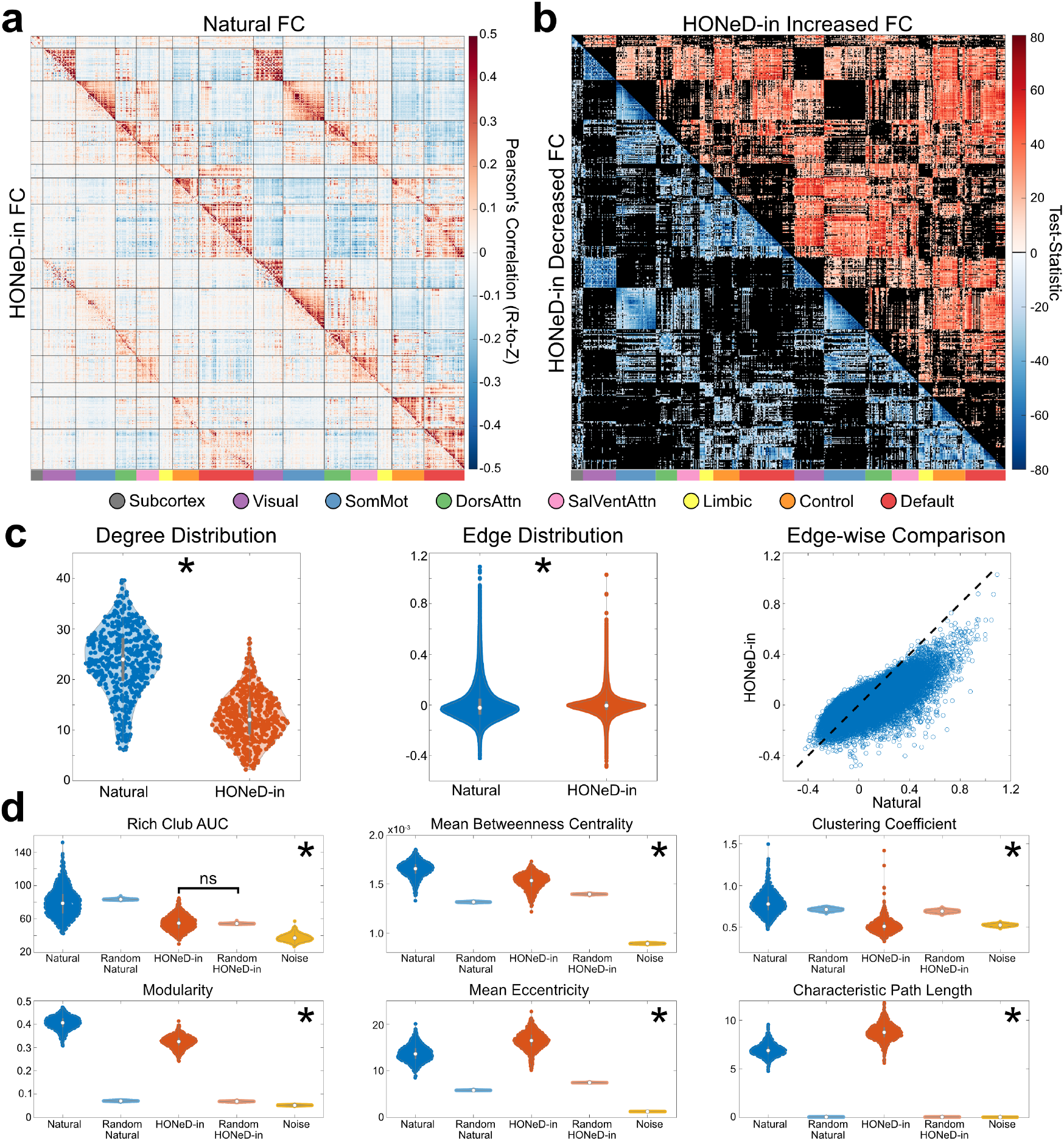
(a) Resting-state FC with the natural FC in the upper-triangular, and the HONeD-in FC in the lowertriangular. Black boarders inside the matrix denote boundaries between canonical RSNs. (b) A summary of mass univariate t-tests on each edge of FC between HONeD-in and Natural. The upper-triangular shows significant increases in HONeD-in FC compared to natural, whereas the lower-triangular shows significant decreases in FC (all Bonferroni corrected). (c) Degree and Edge distribution statistics between the two networks. (d) Six graph metrics to compare natural versus HONeD-in FC as well as randomized networks and a noise network. (*: pairwise comparisons are significant after Bonferroni correction; ns: not significant.)

At the subject-level, we evaluated six common graph metrics to assess the group-level changes in network topology. We found that all six graph metrics were highly significantly changed between natural and HONeD-in FCs (Bonferroni corrected). In general, integrative graph metrics including rich club AUC, betweenness centrality, and clustering coefficient decreased while two segregative measures, eccentricity and characteristic path length, increased. Modularity, although typically considered a segregative measure, also decreased. All measures were also highly significantly different from both randomized networks and the noise network.^9^ Overall, these results highlight how both integration and segregation is altered in HONeD-in FC, with reduced hub-based organization (lower rich club AUC and betweenness centrality), indicating less centralized information routing compared to natural FC.

### HONeD-in FC Changes Resting-State Networks

Since HONeD deconvolution significantly remodels FC, especially pronounced within and between RSNs, we next sought to understand the new RSN structure in HONeD-in FC.

We found empirical RSNs using the original method described by Yeo et al. [74], with von Mises-Fisher (VMF) clustering on both the natural and HONeD-in consensus (group averaged) FC and assessing the RSN organization as well as the clustering stability as also previously defined [74]. We found that 7 networks indeed offered the optimal clustering stability for both natural and HONeD-in RSNs (Figure 4d). However, HONeD-in also had equally good stability with eight network clusters (Supplemental Figure 9). At 7 networks, HONeD-in FC had slightly worse clustering accuracy than natural FC (Natural = 81% versus HONeD-in = 76%, *p <* 10^−36^).

**Figure 4:**
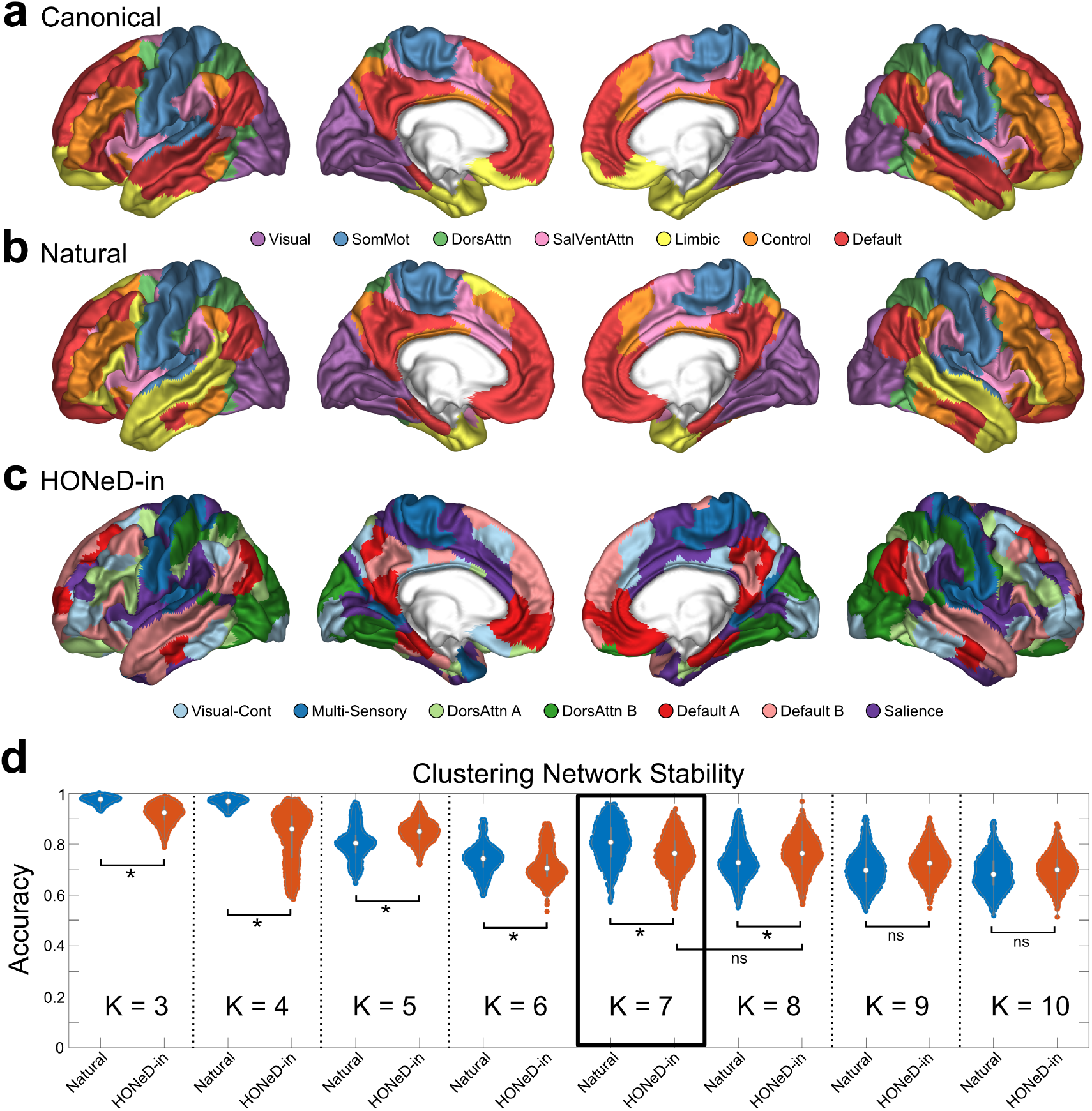
Clustering RSNs. (a) The canonical RSN definition provided by the Schaefer-500 atlas for reference, (b) the most common (mode) clustering assignment for the natural FC, and (c) the most common assignment for the HONeD-in FC. RSNs in the HONeD-in FC have a higher spatial frequency, with networks composed of semi-isolated regions from across the brain. (d) The accuracy across clustering initializations for each of the *K* clusters. Overall, HONeD-in RSNs are slightly but significantly less stable than natural RSNs. HONeD-in RSNs were equally stable with either 7 or 8 clusters. For RSNs plotted separately, see Supplemental Figure 10. For the 8-network HONeD-in clustering, see Supplemental Figure 9. (*: Bonferroni corrected significance; ns: not significant)

The natural FC computed from our empirical data was similar to the canonical RSN definitions, except for the Limbic network, which was temporal-lobe dominant in our clustering result (Figure 4a & b). Meanwhile, the HONeD-in RSNs were starkly different from the canonical organization. First, the Default Mode (DMN) and Dorsal Attention (DAN) networks appear to fracture into two neighboring but distinct functional networks (Figure 4c, DMN: Red & Pink, DAN: Light & Dark Green), both of which show remarkable similarity to previous work reporting that these networks decouple with highly sampled data [9]. We’ve labeled these networks DMN-A/B and DAN-A/B accordingly to align with this prior work. Next, the salience network remains intact in its core areas, however it has also expanded to invade previous somatomotor areas as well as more of the cingulate midline, dorsolateral prefrontal cortex, and both temporal poles (Figure 4c, Purple). The control network merges with primary visual areas, suggesting the formation of a new “visual-control” network (Figure 4c, Light Blue). The previous somatomotor network still partially resides in its canonical location, but now also includes early visual and auditory areas, transforming it into a “multi-sensory” network (Figure 4c, Dark Blue). (For HONeD-in RSN visualizations on separate brain surfaces, see Supplemental Figure 10). When HONeD-in FC has 8 clusters, all RSNs remain the same except for the primary visual regions along the calcarine fissure, which becomes its own 8th network (Supplemental Figure 9).

Overall, the HONeD-in RSNs are fractured and reorganized versions of the canonical RSN structure. This significant remodeling suggests that network diffusion over structural connectivity significantly contributes to the RSN organization in fMRI. Moreover, the changes in these networks indicate that HONeD-in RSNs merge together traditional unimodal regions with multimodal regions.

### HONeD-in RSNs Reorganize Around New FC-Gradients

Given the significant changes HONeD deconvolution imposes on RSN structure, especially the merging together of unimodal with multimodal regions, we next investigated whether these new HONeD-in RSNs had a different relationship to HONeD-in FC-gradients.

First, we assessed how HONeD deconvolution changes the FC-gradients themselves. While the natural consensus FC perfectly reproduces the original FC-gradients, the HONeD-in consensus FC gradients are restructured (Figure 5a). These HONeD-in gradients appear to have high spatial frequency and are distributed throughout the brain with the first gradient qualitatively separating the DMN from Salience regions. Interestingly, HONeD-in gradient 2 has some similarity to the natural gradient 1, except regions are more heterogeneous in the HONeD-in gradient 2 and visual regions decouple.

**Figure 5:**
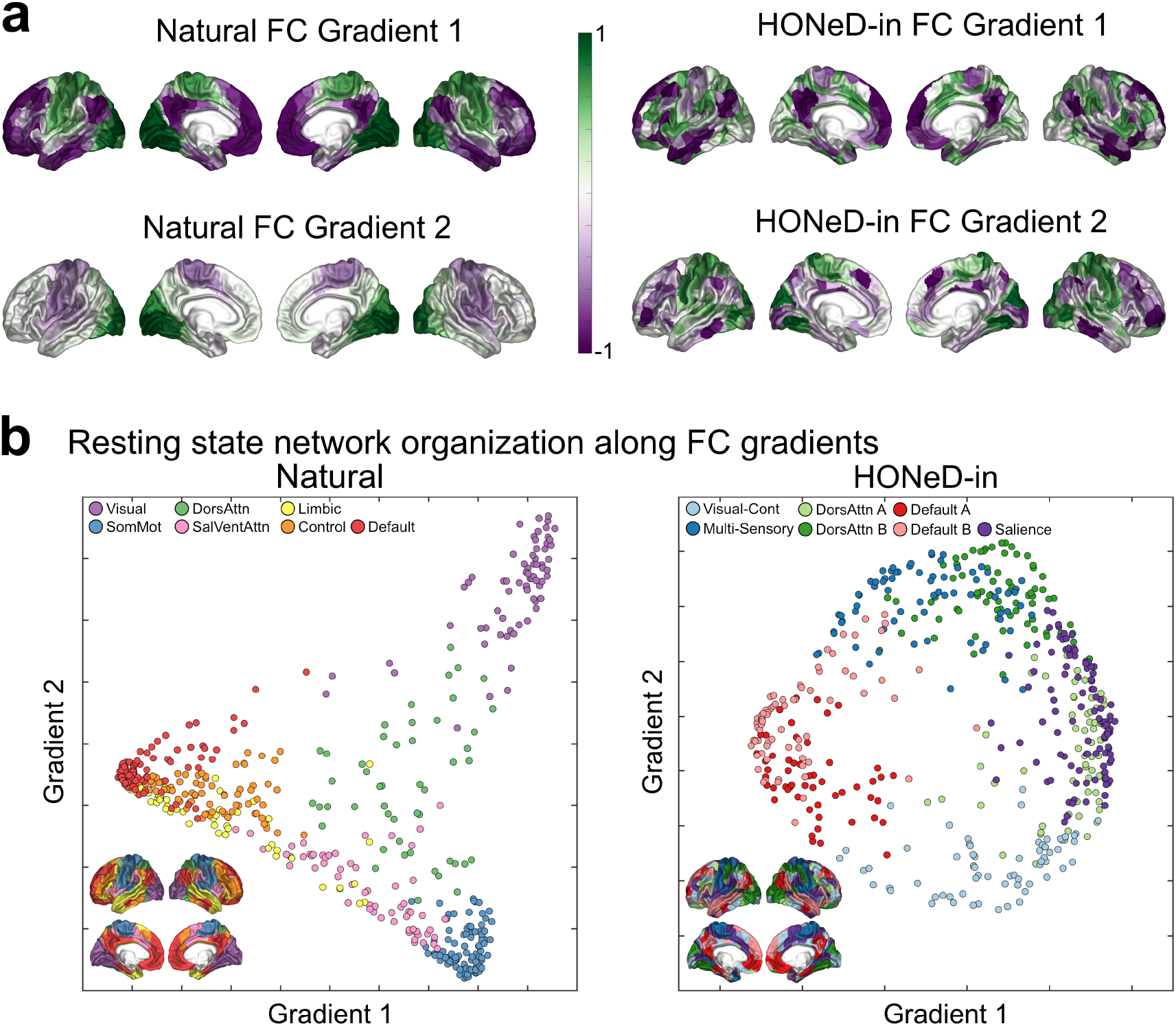
(a) The FC-gradients from natural (left) and HONeD-in (right) signals. (b) RSNs mapped onto FC-gradients derived from the natural FC (left) and HONeD-in FC (right). While the natural FC-gradients place RSNs within a hierarchy organized by a triangle, the HONeD-in FC-gradients instead map RSNs around a circle. The RSN insets are reproduced from Figure 4 for reference.

To better understand FC-gradient relationships to RSNs, we plotted each brain region in the two-dimensional space spanned by the first two FC-gradients, and colored each region by its assigned RSN. The natural FC, as expected, reproduces the triangular hierarchy from previous work [44], with the DMN at one extreme of the first FC-gradient, and the visual and somatomotor networks on the other points of a triangle (Figure 5b left). Strikingly, HONeD-in FC-gradients reorganized RSNs approximately around a circle, thus appearing to deconstruct the previous unimodal– multimodal RSN hierarchy (Figure 5b right). Within the subspace defined by HONeD-in gradients, the DMN networks sit opposed to the Salience network, and the DAN and Multi-Sensory networks sit opposed to the Visual-Control network. Together, these results provide a second line of evidence that the previously-understood hierarchy of RSNs may arise from signal diffusion through the structural connectome, and deconvolving this diffusion dismantles the prior RSN stratification.

### HONeD-in Deblurs Regional Task Activity

We first assessed whether HONeD-in signals had a different hemodynamic response function (HRF) compared to natural signals. We indeed found that the HONeD-in HRF had a stronger peak and earlier decay compared to the natural HRF, but the time to peak was unchanged (Supplemental Figure 11). In the following GLM analyses, we used each signal’s respective HRF set.

We expected that HONeD deconvolution could be useful in increasing the activity of task-related regions while reducing the strength of signal in task-unrelated regions. However, since much of what we know about different cognitive tasks originates with fMRI, motor and language tasks represent the useful benchmarks given their well known regions of interest (ROIs) that are studied outside of fMRI through electrocorticography (ECoG) [42, 52, 66, 65].

In the right-hand motor task contrast, we observed that the natural signal power showed a strongly increased response in the hand regions of precentral gyrus (primary motor cortex) as well as the postcentral gyrus (somatosensory cortex), extending through the inferior parietal lobe (Figure 6a). The effect of deconvolution is especially apparent in this contrast, since the strength of activity in the parietal cortex and postcentral gyrus decreases in some ROIs and eliminated entirely in others; yet, the strength in the primary motor cortex remains. Interestingly, the HONeD-in contrast also strengthens activity in the ipsilateral motor area (Figure 6a), which has been shown to activate during similar motor tasks in ECoG literature [42, 52].

**Figure 6:**
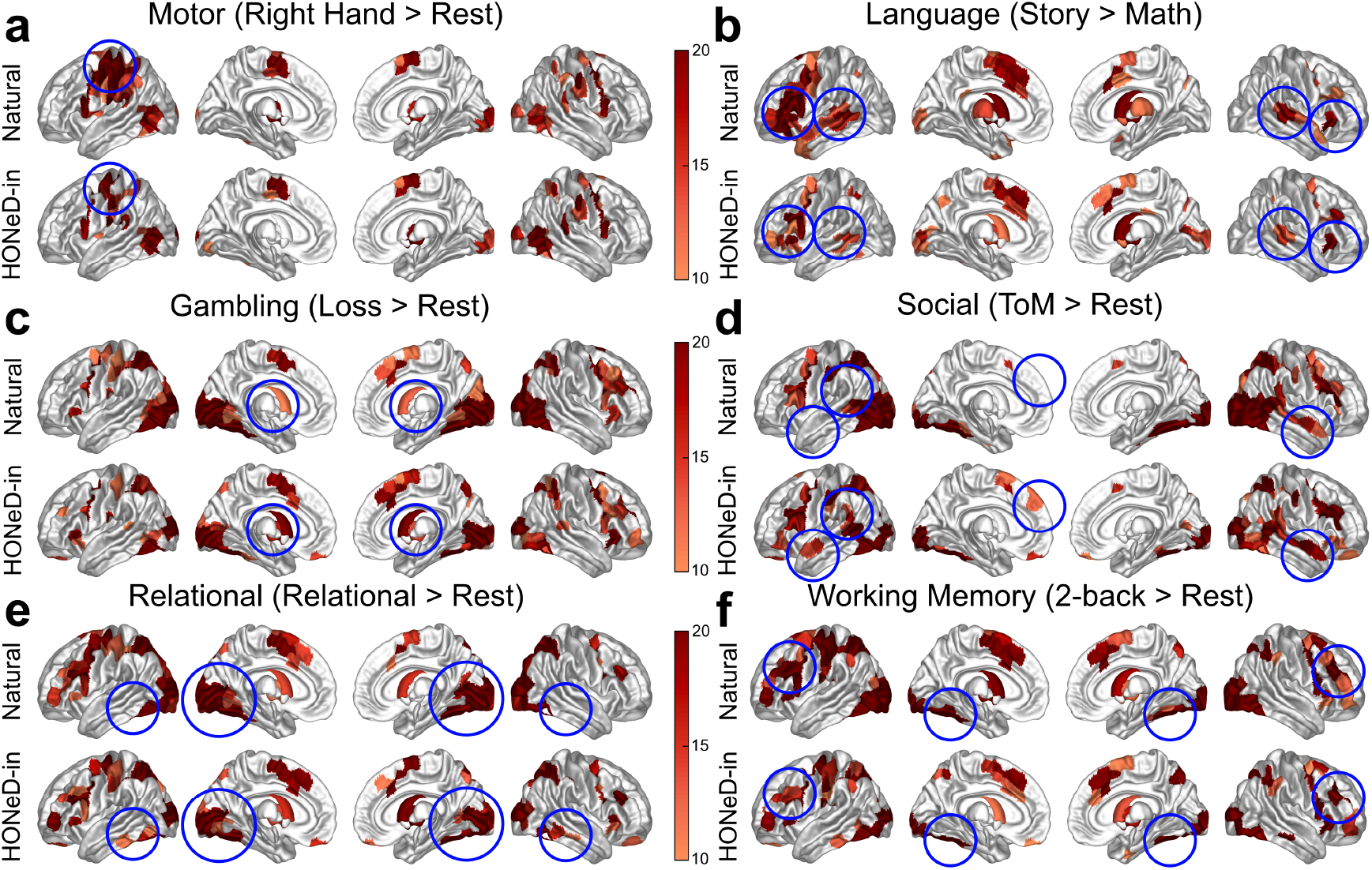
The GLM task contrast for six Human Connectome Project tasks. Task contrasts include (a) Motor (Right Hand *>* Rest) (b) Language (Story *>* Math) (c) Gambling (Loss *>* Rest) (d) Social (Theory of Mind *>* Rest), (e) Relational (Relational *>* Rest), and (f) Working Memory (2-back *>* Rest). In general, HONeD deconvolution appears to “deblur” the activation maps across tasks. For example, the motor cortex activity becomes more restricted to right hand and finger areas during the right finger-tapping motor task (a). Similarly for the language task (b), the HONeD-in signal refines activity around the inferior frontal gyri (Broca’s area) and in well-known language areas in the temporal lobe. The ability for HONeD deconvolution to elevate specificity in activated regions across tasks is nicely portrayed by activity in the primary visual area (V1), where many of the tasks display a significant activity throughout the visual cortex, but the HONeD-in signal tends to refine this signal to V1, with expanded activity during especially visually demanding tasks–Gambling (c), Relational (e) and Working Memory (f). General areas of interesting differences are circled in blue. The color bar is in terms of the test-statistic from the group-level analysis under a Student’s t-test.

Language is another task where we have strong *a priori* ROIs from ECoG, such as the superior posterior temporal sulcus, the middle temporal gyrus, and the inferior frontal gyrus [66, 65]^10^. In the natural contrast, there is high activation in these language areas as well as throughout the temporal lobe (Figure 6b). The HONeD-in contrast eliminates much of this widespread activity while retaining significant activity in regions aligned with previous ECoG literature [66, 65]. Interestingly, HONeD-in contrasts also preserve activity in the right hemisphere, suggestive of bihemispheric language recruitment (Figure 6b).

While the other tasks are more challenging to interpret given their lack of ECoG as a reference, some overarching patterns persist. In general, the HONeD-in contrast represents a sparser and more spatially distributed subset of regions compared to the natural signal contrasts. Some of the regions retained in the HONeD-in contrast are suggestive: the right hand primary motor cortex (relevant to responding to each task) and primary visual areas (relevant to visually relevant tasks). Some qualitatively interesting differences include increased response in reward areas, the caudate and putamen, during the Gambling task, the appearance of theory of mind areas, the dorsomedial prefrontal cortex, right angular gyrus, and anterior temporal lobe, during the social task, and refinements in activity in ventral visual (“what”) areas during Relational and Working Memory tasks (areas circled in Figure 6c-f). However, given the qualitative nature of this analysis without external reference, we refrain from too much speculation about these changes.

## Discussion

Disentangling the role of passive signal diffusion from driving signals in the brain is critical to interpreting macroscale neuroimaging recordings. Here, we developed a method to estimate such a driving signal in closed-form using a higher-order network diffusion (HONeD) model over the structural connectome (SC). The resulting “innovation” (HONeD-in) fMRI signals had significantly altered correlational and hierarchical properties. Specifically, HONeD-in signals 1) sparsified functional connectivity (FC), 2) decoupled resting-state networks (RSNs), 3) reorganized FC-gradients along with the RSN hierarchy, and 4) sharpened task fMRI activation maps. Together, these results suggest that some of the conventional FC structure may be an epiphenomenon of large-scale signal diffusion through SC.

### Passive signal diffusion through SC may confound fMRI

FC from fMRI and SC from dMRI are commonly treated as fundamentally distinct and challenging to reconcile; yet, recent structure-function modeling efforts have made great progress in understanding how FC may arise from SC [22]. Since FC is a statistical measure of covariance between functional time series across regions, SC-FC models are motivated by the hope of endowing FC with biophysical interpretations related to classical neuronal signal propagation along SC. These efforts are especially salient in the active research on complex and nonlinear neural mass models in fMRI [58, 35, 54, 19]. However, the surprising success of simple linear SC-FC models at the macroscopic scale [48], such as the network diffusion-based models [1, 2, 53, 71], suggests that the typical measurements of FC may be confounded by passive signal diffusion over SC. Therefore, HONeD deconvolution seeks to recover an FC that reflects an ‘active’ innovation signal covariance. Although there is no ground-truth to assess our method against, our simulation analyses strongly support HONeD deconvolution as an approach that can recover an underlying structured innovation signal blurred out by passive diffusion over SC (Figure 2). This provides a principled basis for interpreting the following empirical changes we observed after HONeD deconvolution.

### HONeD deconvolution restructures FC and the unimodal–multimodal divide

A pronounced effect of HONeD deconvolution was a global sparsification of FC. HONeD-in FC showed decreased centrality metrics, including Rich Club and betweenness centrality, alongside decreased modularity and increased eccentricity (Figure 3d), suggesting a less hub-dominated and less hierarchically centralized topology. The high network density of FC has been a known challenge in the field of resting-state fMRI connectivity, and approaches to address this issue have included global signal regression (GSR) [41], percolation thresholds [7, 46], or regularization [77]. However, HONeD deconvolution’s advantage over these alternative approaches is that the sparsification arises from a biophysical description of the signal that is being eliminated, rather than a statistical description (e.g. GSR) or a mere pruning of weak edges, which could themselves be meaningful.

Consistent with this global reduction in signal covariance, we found that some of the traditional RSN structure breaks apart. Of particular note were the default mode (DMN) and dorsal attention (DAN) networks, each decoupling into two interdigitated networks (Figure 4). The newly fractured DMN-A was generally posterior and inferior to DMN-B. Meanwhile, DAN-A occupied more frontal areas while DAN-B involved occipital areas, with both interlocking in the superior parietal cortex. We borrowed these A/B division labels from previous work showing remarkably similar RSN decoupling when deeply sampling individual subjects [9, 10]. One difference from this previous work is that we did not observe a specific language network–instead, language areas belonged to DMN-B, which may be a result of our relatively large parcels and that the language network and DMN-B are intermixed through the temporal lobe [10]. Together, our analysis suggests that diffusion through SC may blur together these parallel networks in a way that can only be decoupled with enough resting-state scan time. HONeD deconvolution may help pull apart these parallel networks, providing a new way to study their relevance in fMRI with relatively less data.

Besides fracturing RSNs, HONeD deconvolution also deconstructed the traditional unimodal and multimodal dichotomy in FC. The salience network invaded somatomotor areas seeming to align with the recently proposed integrate-isolate model of the motor homunculus [26], which suggests that the salience network may mediate their proposed action and body representations in a fitting alignment to its role in interoception [40]. The control network reorganizing to include primary visual regions has intriguing parallels with recent work showing a close relationship between control and visual areas during working memory [47]. Instead of a strictly somatomotor network, a new “multi-sensory” network emerged, joining together somatomotor, auditory, and peripheral visual regions (Figure 4c). More than just merging unimodal and multimodal regions in RSNs, HONeD deconvolution in fact dismantled the previous unimodal to multimodal hierarchy along FC-gradients [44]. Instead, our new FC-gradients placed RSNs around a circle, with DMN opposed to the salience and DAN-A networks, and with the visual-control network opposed to the multi-sensory and DAN-B networks (Figure 5b). Together, these findings suggest that dominant network phenomena in fMRI, including RSN organization and unimodal–multimodal dichotomies, may be deeply related to the brain’s SC, highlighting HONeD deconvolution’s potential to investigate these phenomena from a new perspective.

### Implications for task fMRI

Analyzing HONeD-in signals could be especially relevant to task fMRI, where the brain is receiving structured input and generating endogenous computations in response to specific tasks. We found that the GLM-based activation maps from the HONeD-in signal were sparser than activation maps from the natural signal, suggesting that HONeD deconvolution helped to “deblur” these activation maps (Figure 6). It has long been understood that fMRI signals during tasks show strong spatial autocorrelation resulting in large clusters of significant activity [23, 18]. One important source of spatial autocorrelation could arise from passive signal diffusion through white matter, thus blurring the signal effects across regions. HONeD deconvolution precisely aims to correct for this blurring effect, and we indeed observe that the HONeD-in signal reduces large clusters of activity to smaller, spatially separated clusters. In general, we found that regions remaining in HONeD-in activation maps aligned well with the electrophysiology literature on finger movement and language processing [42, 52, 66, 65]. Importantly, HONeD deconvolution did not just sparsify activation maps, but also elevated the activity of important regions. One example was during the theory of mind task where HONeD-in activity increased in the dorsomedial prefrontal cortex and anterior temporal cortex, which are both known to be strongly related to theory of mind [70, 64]. Given how HONeD deconvolution reweights the importance of different regions during tasks, it may provide a useful tool for analyzing task fMRI and localizing task-relevant brain regions.

### Relationship to prior approaches

Several approaches in computational neuroscience share conceptual similarities with HONeD deconvolution. Independent component analysis (ICA) to perform blind source separation was among the earliest methods to search for independent sources of brain activity [12]. Dynamic causal modeling (DCM) takes a different perspective, aiming to find effective (directed) regional relationships in the brain that depend on differences between time-steps [24]. Another approach aims to enhance high-frequency features in fMRI data by isolating the bottom “minor” components of task fMRI time series, thereby creating a “connectome caricature” that improves fingerprinting accuracy [55]. These approaches, however, are statistical in nature and rely solely on the fMRI data without explicit reference to the underlying SC. Network control theory overcomes this limitation by incorporating SC directly to quantify the controllability of different regions and transitions between brain states, and it leverages the same mathematical framework that underlies our derivation of the HONeD-in signal [31, 38]. Another SC-based approach uses graph filtering to extract the high-frequency eigenvectors of the SC Laplacian [28], analogous to the connectome caricature, but instead emphasizing high-frequency *structural* components. HONeD deconvolution nevertheless stands out among these approaches because it provides a closed-form, physics-informed inversion of diffusion on the SC, directly solving for an fMRI innovation signal. In this light, HONeD deconvolution is analogous to the residual of an “interaction picture” in physics, where a simple and well-understood part of the system is removed to reveal more complex dynamics [43]. What remains after HONeD deconvolution is not necessarily a signal untethered from structure: HONeD-in signals isolate the part of the signal that *cannot be predicted by passive diffusion* over the SC, but whose *subsequent propagation* can still be mediated by that same connectome.

### Limitations and future directions

While our HONeD deconvolution method shows promise, there are limitations to consider. One possible concern is that HONeD deconvolution rests in part on obtaining an accurate structural connectome. There are many known limitations with estimating white matter connectivity [34], yet we believe this is not as significant of a limitation as it may first appear. We showed that a consensus SC recovers the innovation signal with as much fidelity as the subject-specific SC, which indicates that many of the highly detailed and subject-specific connections that may be challenging to estimate are not as relevant to HONeD deconvolution. While small variations in SC may not be a significant challenge, the number of brain regions defined for SC and fMRI may prove to be more challenging. We used the Schaefer-500 atlas since it was high-resolution, but still computationally tractable. Task fMRI typically analyzes regional activity at a voxel or vertex level, so extending HONeD to high resolution data would be an important future direction. Another theoretical limitation is that HONeD in its present form assumes that the innovation signal is not correlated across harmonics. However, while harmonics are orthogonal in space, they may not be independent from each other in time. Correlations between harmonics would erode the ability for HONeD-in signals to capture the true innovation signal. This could also be related to the fact that HONeD-in signals include non-uniform actuation arising from a *B* matrix in the LTI equation. Using the HONeD-in signal estimates to recover such a *B* could be an interesting area for future work.

## Conclusion

We introduced a higher-order network diffusion (HONeD) deconvolution to remove the effect of passive signal spread through structural connectivity from functional MRI. This HONeD deconvolution reveals an underlying “innovation” (or “HONeD-in”) signal that substantially remodels fMRI signals. The HONeD-in signal has sparser but still well-connected FC, reorganizes restingstate networks, mixes unimodal and multimodal regions, and deblurs task-evoked activity. Overall, HONeD deconvolution shows significant promise as a new approach to studying cognitive neuroscience and diseases with functional MRI.

## Supplemental Material

### Section 1: Deriving the HONeD-innovation Signal

The solution to our LTI system inhomogeneous ODE is:

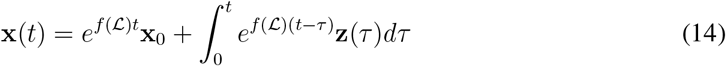

Recall that our higher-order network diffusion (HONeD) model is represented by *f* (ℒ), corresponding to the brain’s internal dynamics. Here, we assume that the innovation signal behaves like a Dirac delta function of the form **z**(*t*) = ∑_*s*_ **z**_*s*_*δ*(*t* − *t*_*s*_) with *t*_*s*_ = *s*Δ*t* such that our sampling interval is Δ*t* = *t*_*s*+1_ − *t*_*s*_ ∀ *s*, which is the signal TR on a uniform sampling grid and where *s* is a dummy index for the summation. Substituting this impulse train into the solution gives the following:

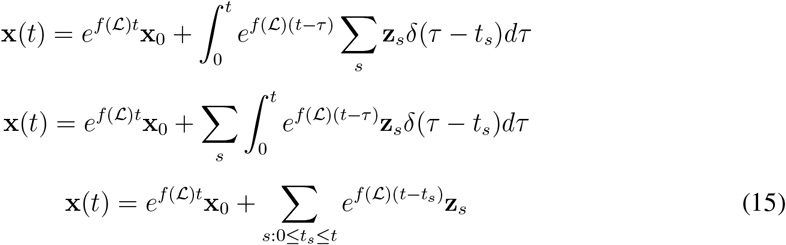

Next, we can write the above equation (15) in discrete time using the following update rule:

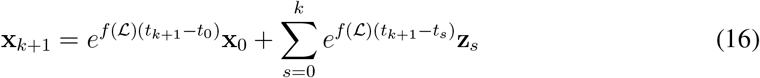

Because an impulse train input **z**(*t*) = _*k*_ **z**_*k*_*δ*(*t* − *t*_*k*_) generally induces jump discontinuities in the state **x**(*t*) at the impulse times {*t*_*k*_}, the value **x**(*t*_*k*_) is ambiguous unless a left/right limit convention is specified. Throughout, we define the sampled state at time *t*_*k*_ to be the left-limit

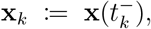

i.e., the state immediately *before* the impulse at *t*_*k*_. Under this convention, the impulse produces an instantaneous jump

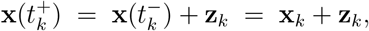

and the state then evolves homogeneously over the interval (*t*_*k*_, *t*_*k*+1_). In other words, we assume that the sampling of **x**_*k*_ happens just before the following innovation signal **z**_*k*_ in the pulse train at time *t*_*k*_. Without loss of generality, we take *t*_0_ = 0. In the following, we will evaluate this update rule for the first two cases and then provide the closed form expression.

Let *k* = 0:

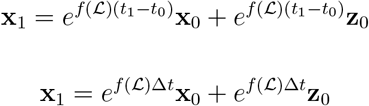

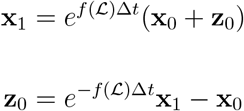

Let *k* = 1:

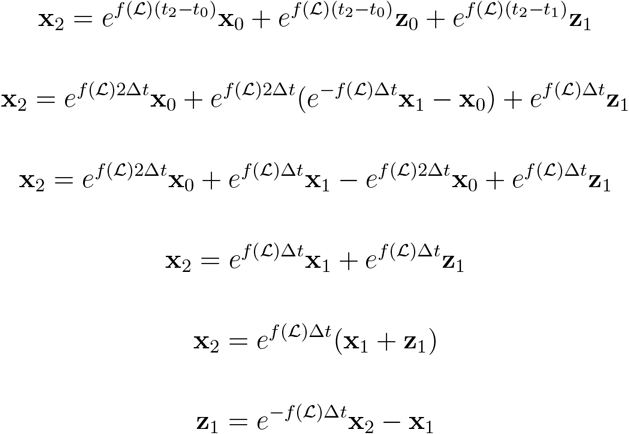

This pattern continues such that the closed form expression for the HONeD-innovation signal at any time *t*_*k*_ is:

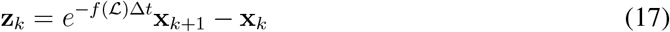

### Extended Data: Figures

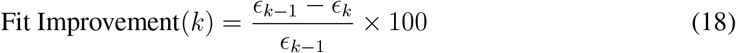

**Figure 7:**
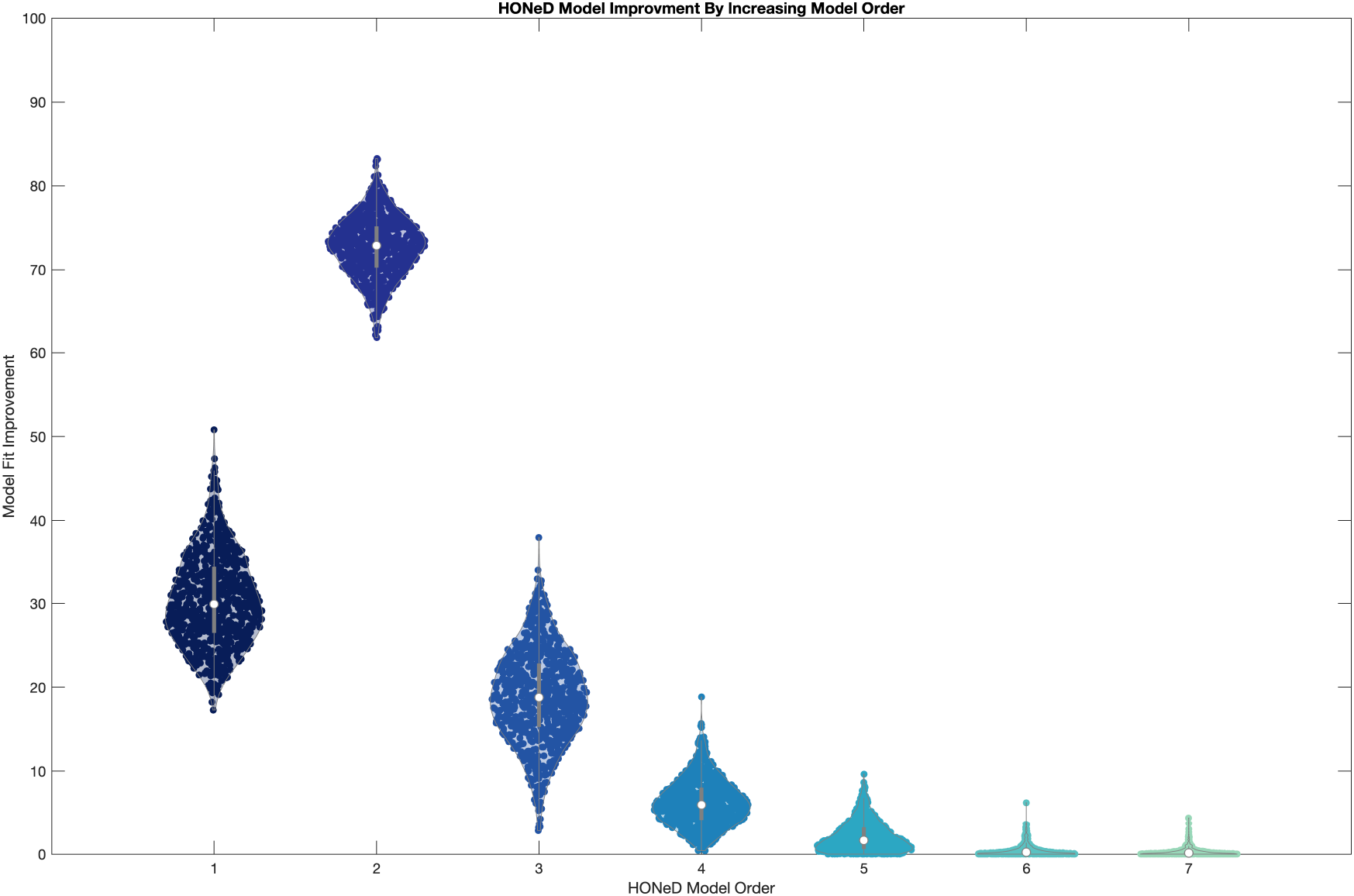
We show the HONeD model fitting improvement in the HCP dataset across varying model orders. Fit improvement is measured in percent and is given by equation 18. The HONeD order = 2 offered the greatest improvement.

**Figure 8:**
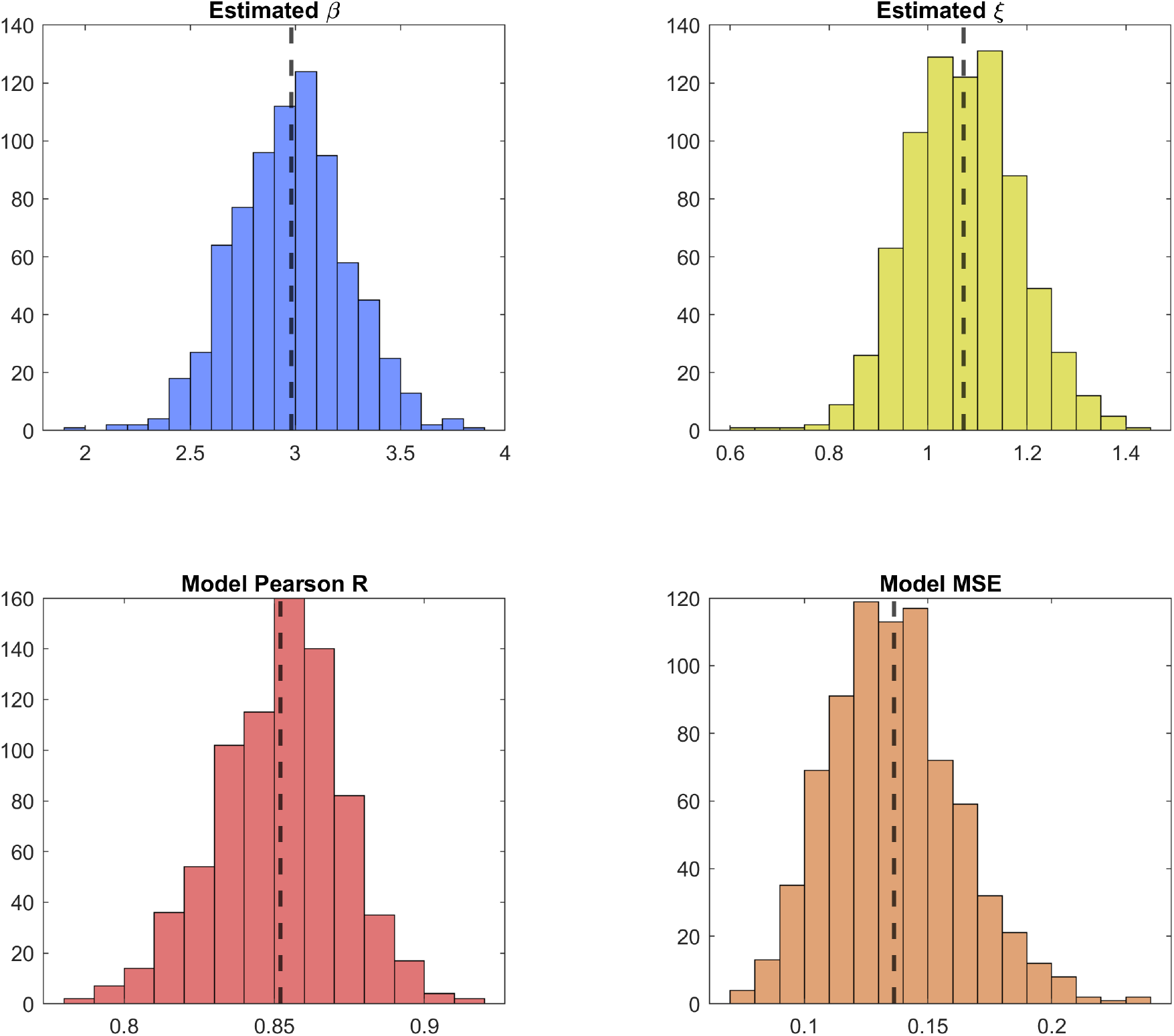
The estimated HONeD parameters, and model performance metrics (Pearson’s R and MSE) between the predicted and empirical graph harmonic power, across the 770 HCP subjects included in this study.

**Figure 9:**
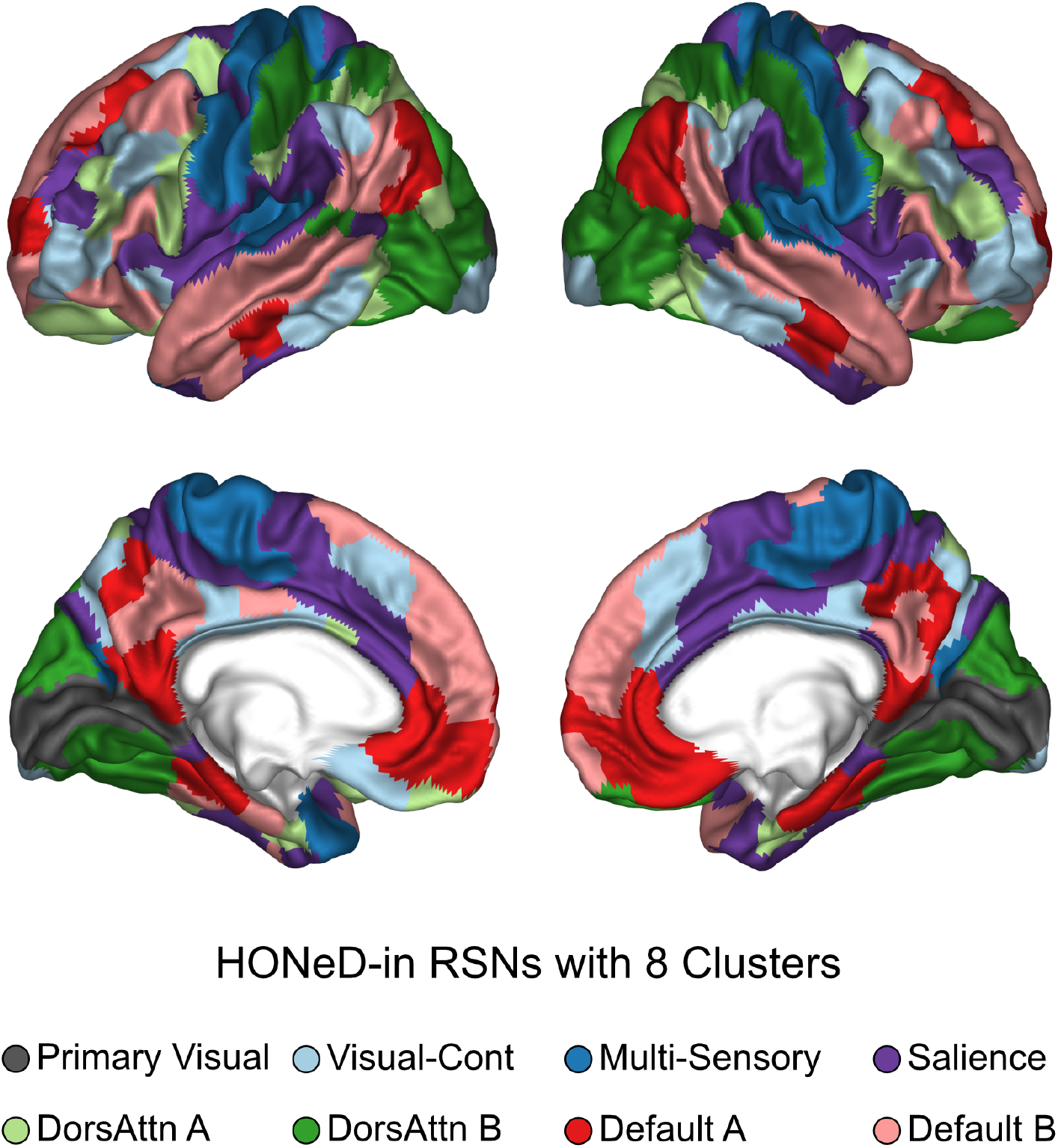
The 8-network solution for von Mises Fisher clustering on HONeD-in FC, which was equally stable to the 7-network solution (shown in Figure 4). RSNs are identical to the 7-network case except for regions around the primary visual areas in the calcarine fissure, which form their own network. Note that the new visual-control network still occupies primary visual areas at the posterior-most end of the visual cortex.

**Figure 10:**
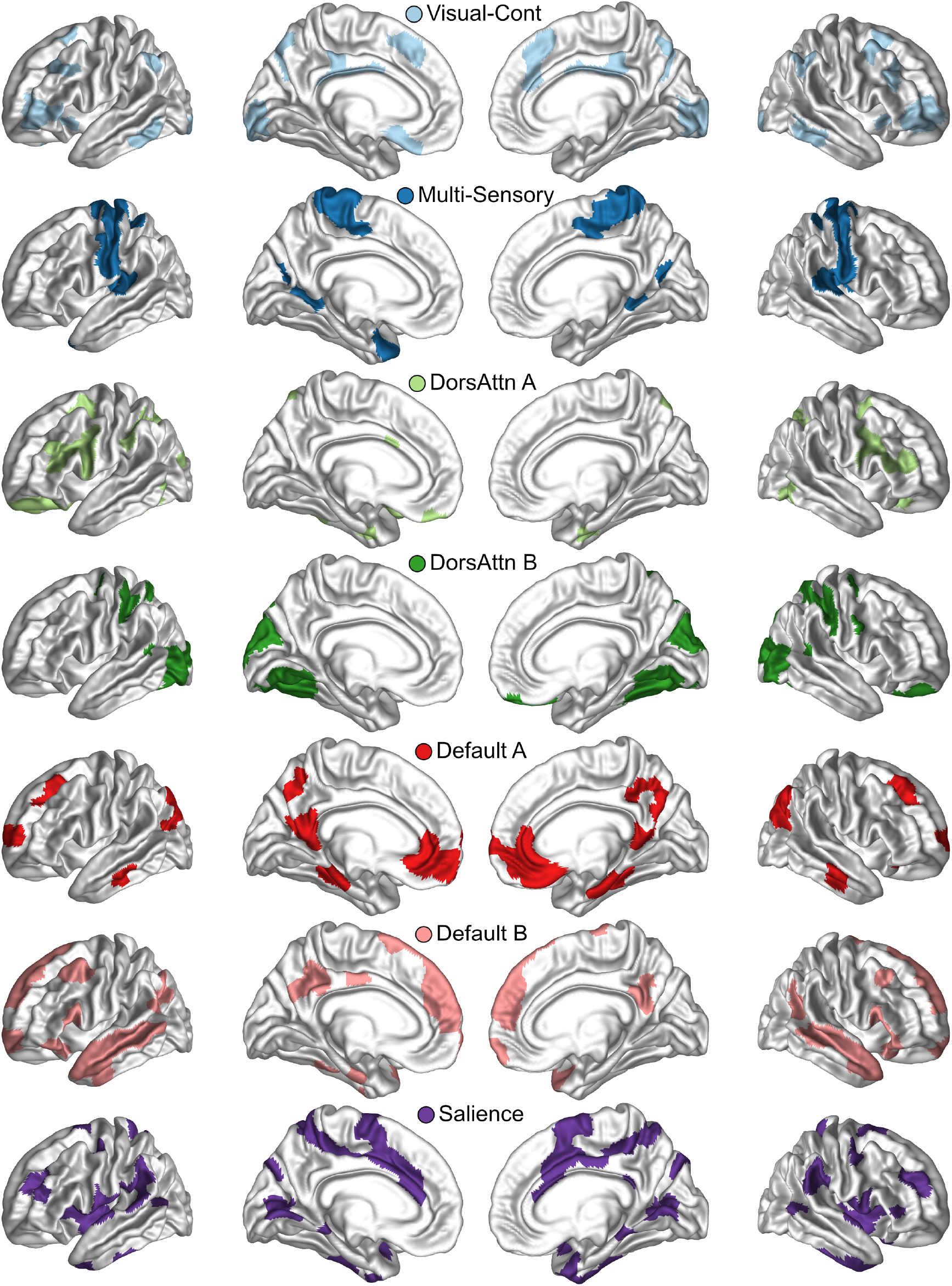
The HONeD-in RSNs shown in Figure 4 plotted on separate brains, with the lateral views rotated 10° for a view of RSN arrangement around gyri.

**Figure 11:**
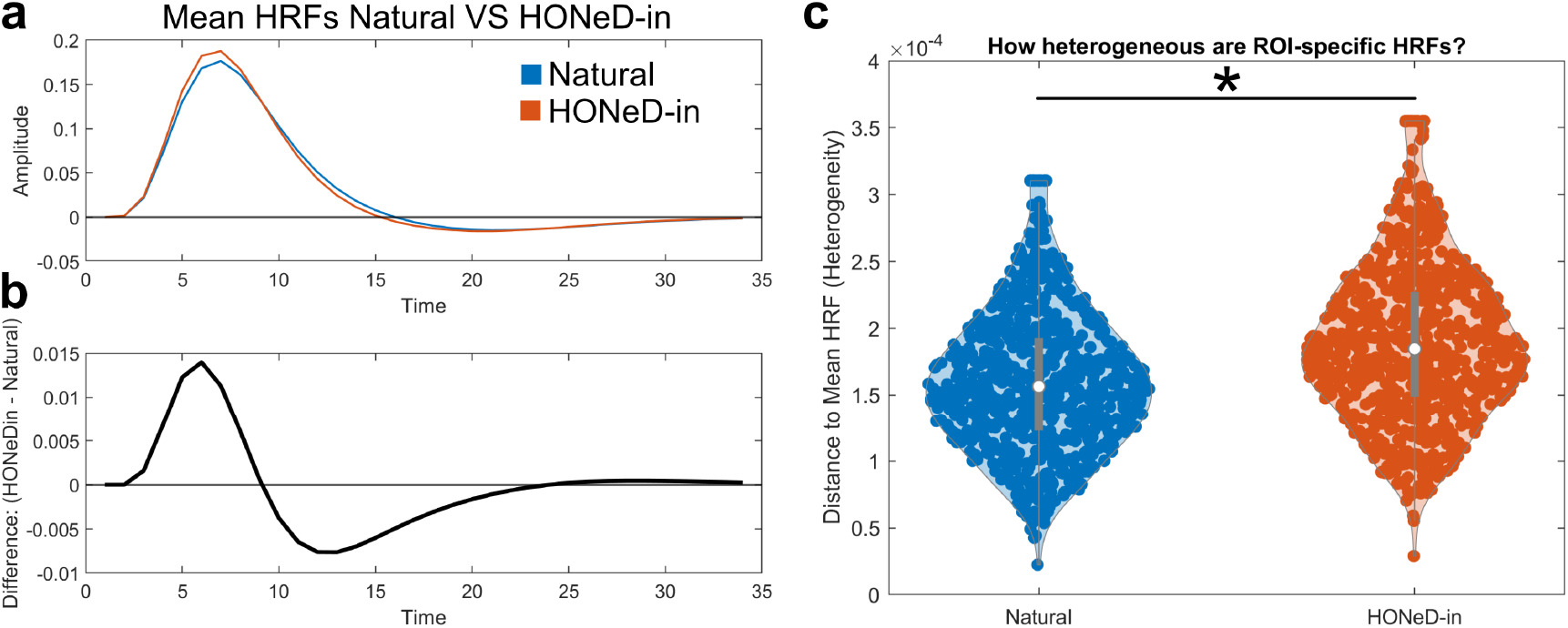
(a) The subject mean estimated hemodynamic response function (HRF) for the natural signal (blue) compared to the HONeD-in signal (orange). (b) The difference between the HONeD-in and natural HRFs, showing that HONeD-in HRF has both a stronger peak and earlier decay compared to natural HRFs. The differences in HRFs were highly statistically significant using cluster-based analysis at all time points (*p*_permutation_ = 0). (c) We evaluated whether region-specific HRFs were more heterogeneous for HONeD-in signals by computing the Euclidean distance between each region-specific HRF and the global mean HRF. HONeD-in HRFs were significantly more heterogeneous than natural HRFs (Student’s t-test: *p <* 10^*−*22^).

Where *ϵ*_*k*_ is the mean squared error for the *k*^*th*^ model order’s fit to harmonic power.

## Acknowledgments

This work was supported by the NIH: R01EB022717, R01AG0621, R01AG072753, R56AG082087, RF1AG062196. Data were provided by the Human Connectome Project, WU-Minn Consortium (Principal Investigators: David Van Essen and Kamil Ugurbil; 1U54MH091657) funded by the 16 NIH Institutes and Centers that support the NIH Blueprint for Neuroscience Research; and by the McDonnell Center for Systems Neuroscience at Washington University

## Footnotes

1 The *p*-th power of any graph’s adjacency matrix, *G*^*p*^ ∀*p* ∈ℕ, computes the movement through the graph after *p*-steps.

2 Note that 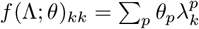 becomes a scalar function.

3 Note that this assumed structure of the innovation signal may not be valid across all neuroimaging modalities.

4 Note that the *homogeneous* diffusion time (*t*) becomes absorbed within the HONeD parameters *θ* .

5 This statistical expectation is because HONeD describes a stochastic processes driven by a random input **x**_0_, meaning we define its theoretical average properties rather than an exact signal. This is justified by the ergodic hypothesis and the assumption that the underlying process is stationary (i.e., statistically stable).

6 Data were provided by the Human Connectome Project, WU-Minn Consortium (Principal Investigators: David Van Essen and Kamil Ugurbil; 1U54MH091657) funded by the 16 NIH Institutes and Centers that support the NIH Blueprint for Neuroscience Research; and by the McDonnell Center for Systems Neuroscience at Washington University. Ethical approval was not required as confirmed by the license attached with the open access data.

7 This global signal is generated by the first eigenmode (*λ*_0_ = 0), which is the ‘DC’ mode and related to the SC’s degree distribution.

8 This ‘*b*’ is unrelated to the diffusion depth parameter (*β*) in the HONeD model.

9 One exception being rich club AUC for deconvolution, which was expected due to the degree-preserving randomization.

10 Historically, the inferior frontal gyrus has been associated with “Broca’s” area, and the posterior superior/middle temporal gyrus has been associated with “Wernicke’s” area, although these divisions are now considered oversimplifications [67].

